# Optimising MALDI on PEN substrates for small molecule analysis in the context of same-section multimodal workflows

**DOI:** 10.64898/2026.09.14.749875

**Authors:** Hugo Delattre, Seren Piper, Ahmed M. A. Abdelhamed, Johanna von Gerichten, Josephine Bunch, Melanie J. Bailey, Rory T. Steven

## Abstract

Multimodal imaging workflows increasingly require multiple analytical techniques to be applied to the same tissue section to improve spatial co-registration and molecular interpretation. Polyethylene naphthalate (PEN) membrane substrates are attractive for this purpose because of their low thickness and compatibility with downstream modalities, but their use for matrix-assisted laser desorption/ionisation mass spectrometry imaging (MALDI MSI) remains challenging due to low conductivity, reduced ion yield and susceptibility to laser-induced damage.

In this study, MALDI MSI conditions for small-molecule analysis on PEN substrates were systematically optimised using bovine brain homogenate and mouse brain tissue using a TimsTOF flex MALDI-2 platform. Matrix selection, laser energy, pixel size (5 and 20 µm), ionisation mode and PEN type were evaluated to identify acquisition conditions that balanced signal intensity with spectral quality and preservation of sample and membrane integrity.

Among the matrices tested, 1,5-diaminonaphthalene (DAN) resulted in the best overall performance, providing higher ion intensities, less damage to the PEN substrates and a broader usable laser energy range than the other matrices tested. Different PEN substrate formats were also compared, showing distinct trade-offs between analytical performance and practical suitability for multimodal analysis. Under the tested conditions, glass-supported PEN (gPEN) provided the most practical overall compromise, while frame-supported PEN was not suitable for routine analysis on the instrument configuration used.

Application of the optimised method to mouse brain tissue confirmed that spatially resolved metabolite imaging on PEN substrates is feasible and can support same-section multimodal workflows.

## Introduction

Matrix-assisted laser desorption/ionisation (MALDI) mass spectrometry imaging (MSI) can be used to map the spatial distributions of a wide range of molecules, including metabolites and other small molecules^1, 2^, from diverse sample types. The distributions of small molecules in biological samples can provide valuable insight into alterations caused by disease, drugs or other factors^3, 4^. Due to the complexity of the composition of tissues, MALDI is increasingly integrated with complementary techniques to generate a more complete biological picture than any single modality alone^2, 5, 6^.

Acquiring data using a single tissue section strengthens multimodal workflows by improving the accuracy of spatial dataset integration and increasing profiling fidelity to specific tissue regions, particularly in heterogenous samples^2, 7-9^. Polymer membranes such as polyethylene naphthalate (PEN) are compatible substrates t for various imaging modalities (*e*.*g*. ion beam analysis (IBA)^10, 11^, X-ray fluorescence (XRF), desorption electrospray ionisation (DESI), secondary ion mass spectrometry (SIMS), stimulated Raman scattering (SRS)^12^, and laser capture microdissection (LCM)^13, 14^) and therefore are sensible candidates for a multi-modal workflow involving MSI. They stand apart from typical MALDI substrates with low conductivity and thickness (typically 1-4 µm) while being composed of organic molecules with low concentrations of trace elements.

Performing MALDI on polymer substrates with ultraviolet (UV) lasers is intrinsically challenging due to their susceptibility to laser damage and low conductivity, which can lead to surface charging effects which, in turn, contribute to a reduction in ion counts. A trade-off must be achieved of sufficient laser power to generate signal for imaging while preserving the sample and membrane for downstream analysis.

The results of a MALDI analysis depend heavily on matrix choice and instrument parameters, particularly for small molecule imaging where ion-suppression effects and poor inherent ionisation efficiency further compromise results^15-17^. Parameters that affect the amount and rate of laser energy transferred to each point on the sample include pixel size, laser power, laser firing frequency and the number of laser shots. MALDI-2 post-ionisation has recently become a promising tool for increasing small molecule intensities by firing a second UV laser perpendicular to the direction of ion extraction^18-20^. Another important consideration is the choice of PEN substrate, since the membranes are commercially available as both glass-backed and metal framed. Without optimisation, integration of MALDI into multimodal workflows may lack robustness due to the number of factors to address.

In this study, we systematically optimised MALDI MSI for small-molecule analysis on PEN membrane substrates in the context of same-section multimodal workflows. Matrix composition, laser power, pixel size, ionisation mode and PEN substrate type were evaluated to identify acquisition conditions that provide sufficient signal while preserving sample and membrane integrity for sequential analysis. This work establishes a practical framework for selecting matrices, substrates and acquisition parameters for MALDI analysis on PEN within multimodal imaging pipelines.

## Materials & Methods

### Chemicals & substrates

Deionised water (ρ =15 MΩcm–1) was produced in-house using an Elga Purelab system. All solvents were purchased from Fisher Scientific (Loughborough, UK) and matrix compounds from Sigma-Aldrich (Merck, Gillingham, UK). Indium tin oxide (ITO; 70-100 Ωm) coated glass slides were purchased from Sigma-Aldrich (Merck, Gillingham, UK). 4 µm thick glass and frame polyethylene naphthalate (gPEN and fPEN) membrane slides were obtained from Leica Microsystems (Milton Keynes, UK).

### Tissue sectioning

Homemade bovine brain homogenate (BBH) and wildtype mouse brain in the coronal orientation were sectioned at 10 µm thickness using a CryoStar NX70 (ThermoFisher; -15 oC blade temperature, -13 oC specimen temperature) and thaw mounted on ITO, gPEN or fPEN slides pre-washed with ethanol. When sectioning onto fPEN slides, the tissue was mounted into the recessed cavity side of the fPEN substrate frame. Samples were dried under nitrogen, vacuum-sealed and stored at −80 °C. All animals and tissue were managed in accordance with the UK Home Office Animals (Scientific Procedures) Act 1986.

### Matrix application

For all samples, a 7 mg/mL solution of matrix was prepared in 90% methanol and 10% water and sonicated for 20 minutes. The matrix solution was applied using a M3+ sprayer (HTX technologies) using the parameters: 60 oC nozzle temperature, 14 passes, 0.074 mL/min flow rate, 1000 mm/min sprayer velocity, 3 track passes.

### MALDI analysis

All datasets were acquired on a TimsTOF Flex microgrid MALDI-2 (Bruker Daltonics) with the global attenuator offset position at 0%. Table S1 contains the instrument parameters used for data acquisition, apart from laser energy. Mass calibrations were performed using red phosphorous pipetted onto the sample slides. Optical images of all samples were taken with an Axio Imager M2 (Zeiss, Cambridge) brightfield microscope before and after MALDI analysis.

### Instrument parameter and matrix comparison

BBH tissue was sectioned onto two gPEN slides, each containing a single tissue section. Each section was divided into two halves, with each half coated with a different matrix: 9-aminoacridine (9AA), 1,5-diaminonaphthalene (DAN), 2,5-dihydroxybenzoic acid (DHB), or N-(1-naphthyl)ethylenediamine dihydrochloride (NEDC).

Within each matrix-coated region, data were acquired at two pixel sizes (5 and 20 µm) and in both MALDI and MALDI-2 modes. For each combination of matrix, pixel size, and ionisation mode, regions of interest (ROIs; 500 pixels) were defined on the tissue and acquired in a randomised order. Laser energy was varied from 10% to 100% in 10% increments across acquisition methods. For MALDI-2 experiments, the instrument automatically applied a 15% increase to the primary laser energy setting.

### Substrate comparison

After sectioning, fPEN samples were mounted onto ITO slides (itoPEN; SI Figure S3) using a minimal amount of cyanoacrylate adhesive applied to the membrane edges, away from the MALDI analysis region. The membrane was pressed onto the ITO slide under a weight for 5 min, after which the surrounding frame was separated from the membrane with a scalpel.

DAN was sprayed onto BBH tissue samples mounted on four different substrates (fPEN, gPEN, itoPEN and ITO). If air was trapped behind the membrane (for itoPEN and gPEN), small holes were made with a scalpel after matrix deposition to enable its release.

Due to size incompatibility with the instrument slide holder (MTP Slide Adaptor II), gPEN slides were resized, using a combination of scratching with a diamond tip pen then snapping off the edge and filing with a metal file. The fPEN slides were mounted onto a MALDI spot plate (MTP 384 target plate ground steel) using copper tape.

Guided by the laser energy optimisation experiments, a reduced range of laser energies were acquired for each combination of pixel size (5 and 20 µm) and ionisation mode.

### Mouse brain imaging

DAN was sprayed onto coronal mouse brain sections on ITO and gPEN substrates. Each slide had two sections, one was analysed with MALDI and the other with MALDI-2, all using 20 µm pixels. Each brain section was split in half into two regions of interest, which were acquired with different laser energies – the OLE and MVLE specific to that combination of pixel size and ionisation mode.

### Cell imaging

PEN slides were sterilised in 70% EtOH for 20 min, dried at RT and kept under UV light for 1 h. Human pancreatic adenocarcinoma cells PANC-1 (Merck, UK) were cultured in DMEM high glucose (Fisher Scientific, UK) with 10 % (v/v) fetal bovine serum (Fisher Scientific), 1 % penicillin/ streptomycin (Fisher Scientific), and 2 mM L-glutamine (Sigma-Aldrich). Cells were kept at 37 °C with 21 % O_2_ and 5 % CO_2_.

150,000 PANC-1 cells (P25) per well were cultured on gPEN using Millicell® EZ Slide 4-well, 1.7 cm^2^/well and incubated for 3 days. Cell culture media and then the Millicell® EZ Slide 4-well were removed and the slide was washed with 37 oC phosphate-buffered saline (PBS) for ∼30 s. Cells were fixed with 4% paraformaldehyde (PFA, Fisher Scientific) for 5 min then washed twice with room temperature PBS followed by deionised water and left to dry for 20 min. The slide was stored at –80 oC until analysis with MALDI-2, 10 µm pixel size and 50% laser energy.

PANC-1 cells were cultured on fPEN in a petri dish and incubated for 2 days until they reached ∼60% confluency. Cell culture media was removed, and the slide was washed with PBS and 0.15 M ammonium formate three times each. The cells were snap-frozen by floating in liquid nitrogen then freeze-dried (FreeZone 2.5L, Labconco) overnight with the condenser at -80oC under vacuum (0.75 mbar). The slide was stored at -80oC until it was made into itoPEN (described above) and analysed with MALDI-2, 10 µm pixel size and 50% laser energy. A second region was analysed at 60% laser energy using *m/z* 500-1100 range.

### MALDI data analysis

The data were analysed in SCiLS (Bruker Daltonics) and MATLAB (MathWorks) software. All datasets were recalibrated to their respective matrix peak in DataAnalysis (Bruker Daltonics) using ‘Linear Correction’ before creating SCiLS files. Regions of interest that excluded background and holes in the tissue homogenates were created prior to calculating the on-tissue ion intensities. Grouped bar graphs were plotted using the median values and the median absolute deviation for error bars.

### Scanning electron microscopy (SEM)

All DAN matrix samples used for laser optimisation were coated with 3 nm of gold using a Q150V ES Plus (Quorum, Laughton, UK) with 99.999% gold sputter target. Micrographs of all acquired regions were acquired on a Apreo 2 SEM (Thermo Fisher Scientific, Waltham, USA) with the Everhart-Thornley detector (ETD) at 800x, 1500x and 2000x magnifications (horizontal field width 159 µm, 84.7 µm and 63.6 µm respectively) with an accelerating voltage of 5.00 kV, beam current of 0.10 nA and working distance ∼10 mm.

## Results & Discussion

### Instrument parameter and matrix comparison

PEN membranes exhibit strong optical absorption in the near-UV region ^21, 22^, meaning that lasers of typical (commercial) MALDI MSI systems will readily ablate holes in the membrane, resulting in poor quality MALDI data and the inability to carry out downstream analyses. Therefore, we evaluated the laser energy (%) parameter as a first step in method development. The relevant instrument parameters are the pixel size, number of laser shots per pixel, laser repetition rate and laser energy (%). To reduce the dimensionality of the optimisation, the laser repetition rate and number of shots were fixed at routinely used values (SI Table S1). The median ion intensities for selected putatively assigned metabolite ions (SI Table S2) across the investigated range of laser energies for BBH on PEN are shown in **Figure 1**.

**Figure 1.**
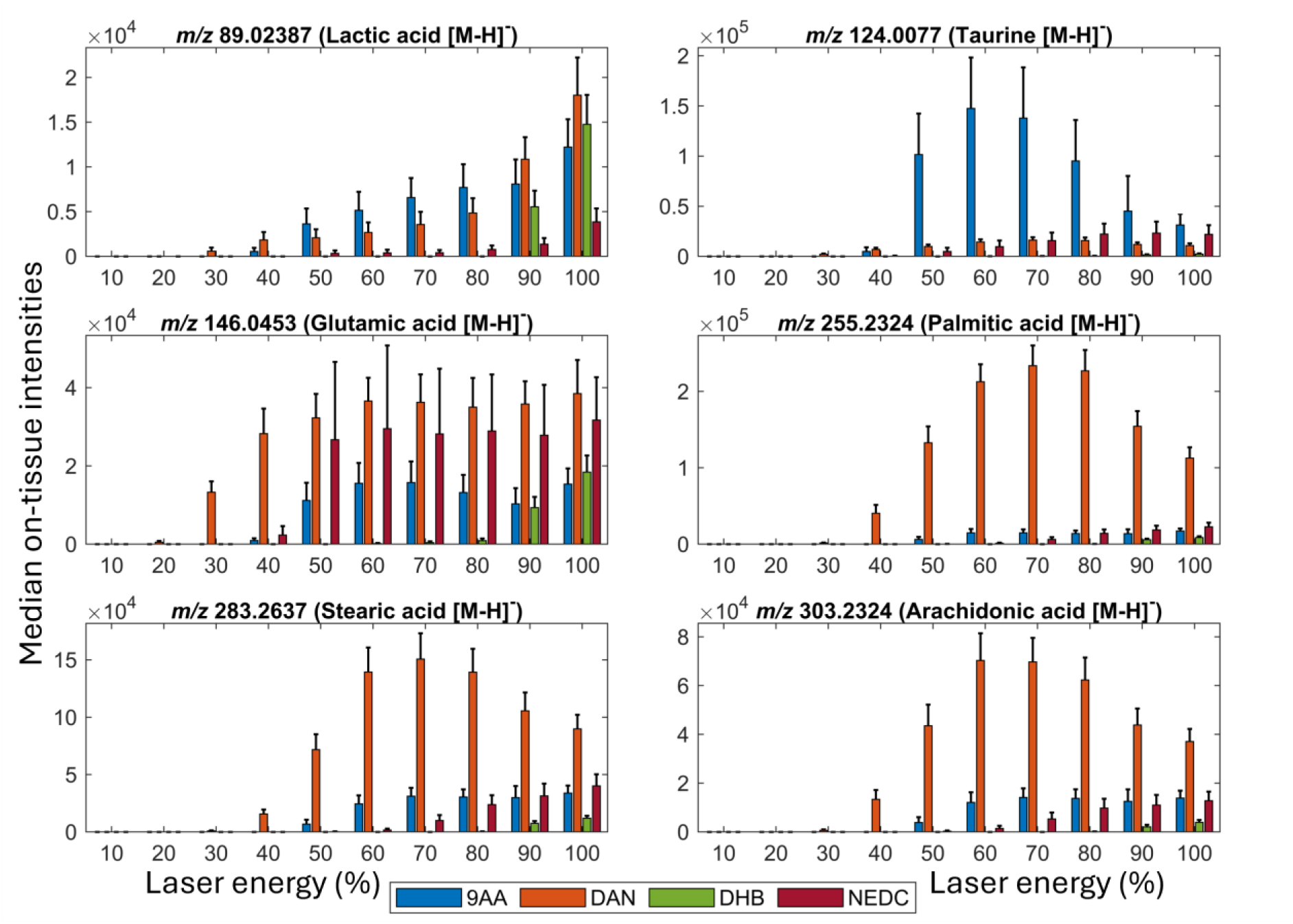
Grouped bar charts of median on-tissue intensities across instrument laser energy settings for deprotonated ions of interest, acquired on BBH tissue mounted on gPEN. Different coloured bars for each matrix are shown at each laser energy. Data acquired with 20 µm pixel data without post-ionisation is shown. Assignments are putative and intensities within ±15 ppm are summed.

Distinct responses were observed for each of the matrices, depending on the laser energy. For most ions, the greatest median intensities were produced by the DAN-coated sample across the laser energy range. Exceptions were observed for lactic acid and taurine, for which 9AA performed better overall. However, at low laser energies between 10 and 40%, DAN still performed better. For lactic acid, DAN-coated tissue also produced the highest intensities at laser energies above 80%, though this may indicate a spurious contribution to ion signal at this *m/z*. 9AA also performed well across the range of laser energies, however detectable ion signals were generally obtained at higher laser energies than DAN, with a particularly strong at response for taurine. The sample coated with NEDC produced detectable intensities at slightly higher laser energies than 9AA, but these were usually lower, except for glutamic acid, which NEDC was effective at ionising. DHB required the highest laser energy settings before producing detectable intensities, and still only produced comparatively low ion counts.

Across all combinations of pixel size and ionisation mode, DAN-coated tissue produced the highest overall intensities (SI Figure S4). Pixel size had less of an effect on the trends seen in Figure 1 than the ionisation mode. With MALDI-2 enabled, the proportional intensity differences between matrices were smaller. Taurine was not detected with DHB as the matrix and MALDI-2 enabled.

The varying laser energy at which matrices can produce detectable ion intensities of metabolites is attributed to a combination of the relative matrix-specific laser fluence thresholds as well as analyte ionisation efficiencies^20, 23-26^. For MALDI integration into multimodal workflows, it is necessary to define suitable working ranges of parameters, such as laser energy. This establishes a balance between detected ion intensities and minimising tissue and substrate damage, such that downstream analysis by additional analytical modalities remains feasible. To this end, the ‘minimum viable laser energy’ (MVLE) was defined, qualitatively, as the lowest laser energy setting that produced useful spectral and imaging data. Ion images from a set of putatively assigned metabolites of interest were visually assessed for tissue-associated spatial distributions. Hole and edge features in tissue homogenate sections were used to make this judgement. From this assessment, the lowest median number of counts required, on tissue, to resolve the tissue holes was found to be approximately 200 for datasets imaged at 5 µm pixel sizes and 500 for 20 µm pixel sizes.

The upper bound of laser energy suitable for analysis of tissue on PEN was defined as the ‘optimal laser energy (OLE)’, corresponding to the lowest laser energy setting that maximised signal across the selected ions of interest, thus not incurring excess damage for little further increase in detected ion intensity. This upper bound is important since, in addition to damaging the PEN membrane, excessive laser power can promote fragmentation of molecules and an overall reduction in signal for an ion^27, 28^. Since the optimal laser energy is analyte-dependent, the OLE was selected as determined as the best compromise across all six ions of interest. Figure 2 compares the MVLE and OLE values across all datasets. MVLE and OLE % values for each matrix are shown in the sub-figure legends.

**Figure 2.**
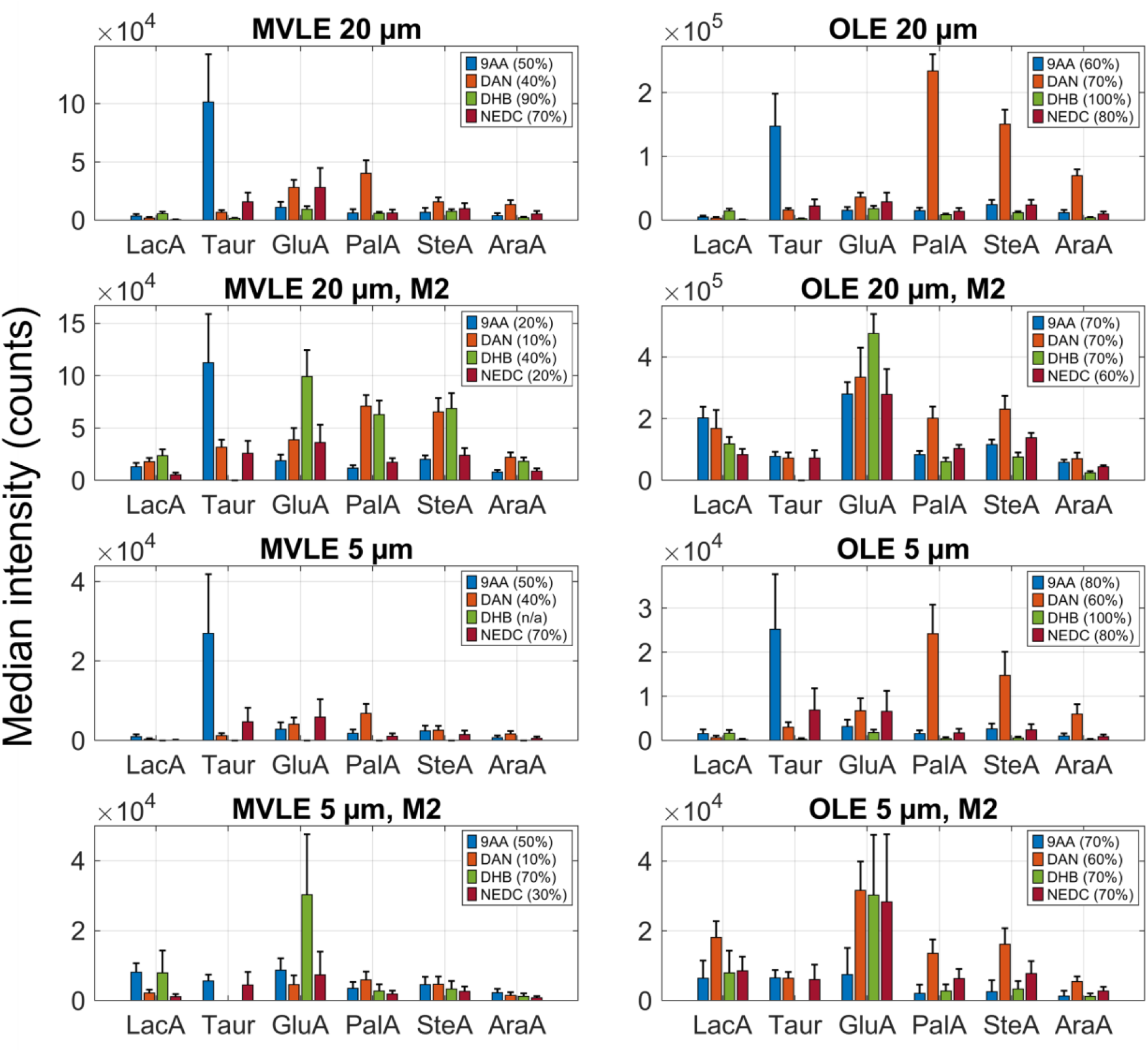
Grouped bar chart comparing the laser energy settings and intensities of all MVLE and OLE for a selection of analytes. The same six ions of interest as in **Figure 1** are putatively assigned and intensities are summed around the peak ±15 ppm. LacA = Lactic acid, Taur = Taurine, GluA = Glutamic acid, PalA = Palmitic acid, SteA = Stearic acid, AraA = Arachidonic acid, M2 = MALDI-2. Only the error bars above the median are shown for clarity.

DAN-coated tissue produced the highest intensities at the OLE and as well as performing well at MVLE was assigned the lowest MVLE % energy in both MALDI and MALDI-2 modes. It was also assigned the widest MVLE to OLE range, indicating greater tolerance to variation in acquisition conditions. This adaptability is ideal for multimodal workflows where MALDI is not the final step and minimising sample damage is important. Overall, DAN was judged to outperform the other matrices and was selected for use in further analysis and experiments. Figure 2 also shows that, even after accounting for the automatic MALDI-2 boost in the primary laser energy (+15%), which was a feature of the Bruker instrument control software “timsControl” at the time, enabling MALDI-2 post-ionisation lowers the MVLE threshold, which may reduce the damage to the polymer substrate. Using MALDI-2 post-ionisation also expanded the laser energy range between the MVLE and OLE, potentially providing greater flexibility when balancing analytical performance against preservation of tissue and substrate integrity.

In addition to review of single ion trends related to laser energy %, overall spectral quality, provides important context. Since the membranes are known to be easily damaged by the laser, they can introduce background peaks of polymer fragments into spectra. **Figure 3** shows the spectra for the MVLE and OLE of all combinations of pixel size and ionisation mode, with enlarged views of the *m/z* regions corresponding to the six ions of interest reported above.

**Figure 3.**
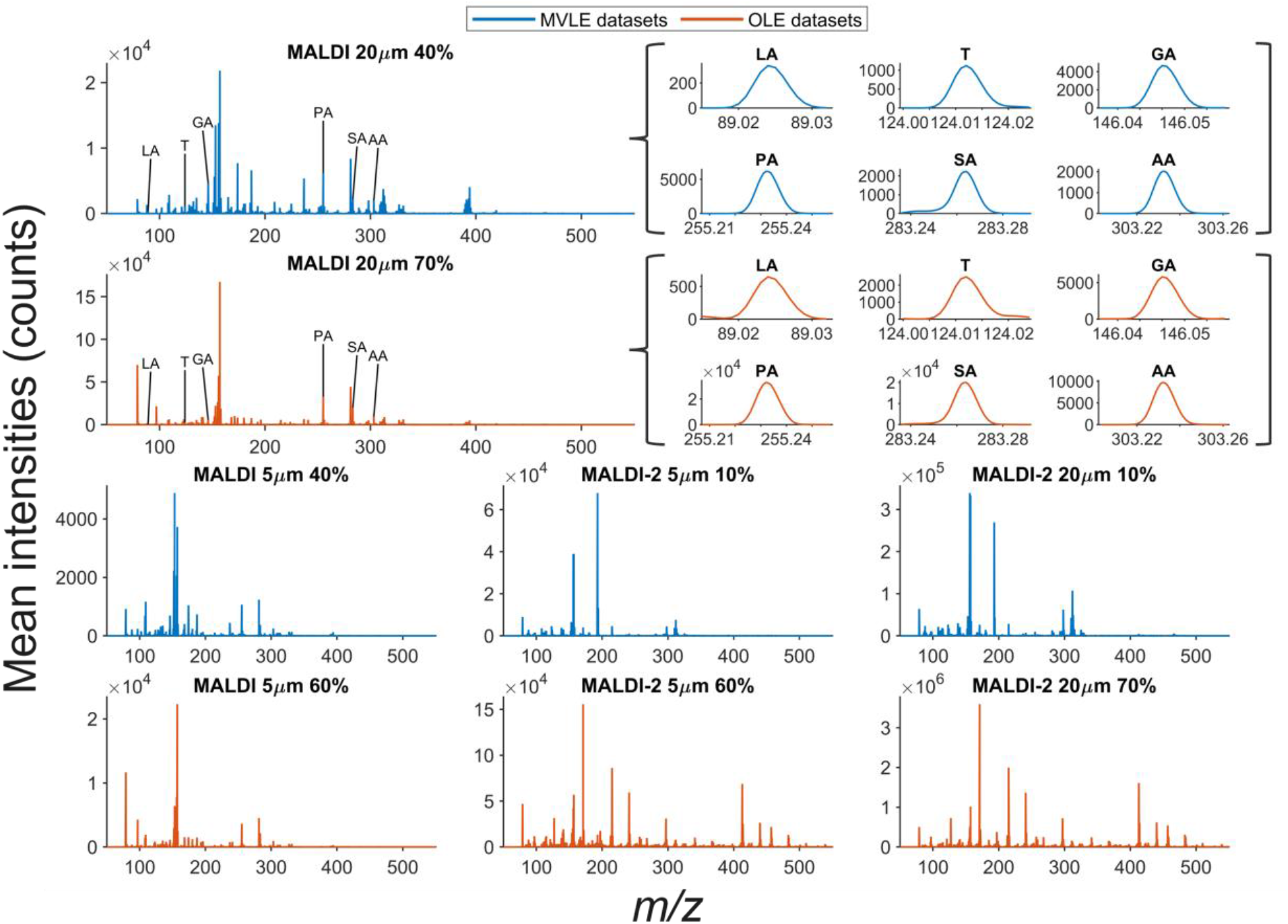
Mean on-tissue spectra from the DAN-coated sample at the MVLE and OLE settings collected at 5 and 20 µm pixel pitch. All the MVLE spectra are plotted with blue traces, while the OLE spectra are plotted with orange. Plots with reduced *m/z* ranges are included for the top two mean spectra to show the peaks putatively assigned to the six ions of interest referenced above.

Increasing the laser energy from the MVLE to OLE did not substantially alter the overall spectral profile, although peak intensities generally increased, demonstrating that analyte detection can be achieved while minimising substrate damage. In datasets acquired without post-ionisation, MVLE and OLE spectra were broadly similar at 5 µm and 20 µm pixel sizes, although peaks above 300 *m/z* were generally less prominent at 5 µm pixel size. As expected, larger differences were observed between datasets with and without MALDI-2 post-ionisation at the same pixel size. MALDI-2 generally increased both the number and intensity of peaks, particularly at the OLE. This likely reflects the combined effects of the automatic boost in the primary laser energy (+15%), in addition to the action of the post-ionisation laser. Furthermore, peaks in the zoomed *m/z* plots indicate that all six metabolite ions remained readily detectable and exhibited well-defined peak shapes across acquisition conditions.

Comparison with reported PEN fragment^29^ *m/z* values identified several possible matches in the spectra (SI Table S5) across all samples. The origin of the additional ion peaks above 300 *m/*z could be from post-ionised neutral species, in-source fragmentation, substrate-derived ions, or a combination of these. Although no off-tissue pixels were available for direct background assessment, the holes in the tissue homogenate provided regions with minimal tissue-endogenous ion contribution. The candidate background peaks showed higher intensity in these holes, while remaining detectable across the entire ROIs, and were present in samples coated with all the other matrices, which argues against matrix peaks as a possible origin. Together, these observations indicate that the additional peaks are likely to represent background signal, potentially arising from the PEN membrane or surface contamination.

As these background peaks did not overlap with the ions of interest, subsequent analysis focused on defining the sample and membrane damage limits. If the laser energy setting is set too high, there may be insufficient sample left for subsequent modalities to produce useful data, which is important if MALDI is not the final modality in the workflow. Even if it is the final modality, it is important that the structural integrity of the substrate is maintained so that the sample retains planarity and, at the extreme, is not completely ablated as with LCM. Figure 4 shows tiled microscope images of the acquired regions on the tissue homogenates for each of the matrices and a scanning electron microscope image panel for DAN coated homogenate.

**Figure 4.**
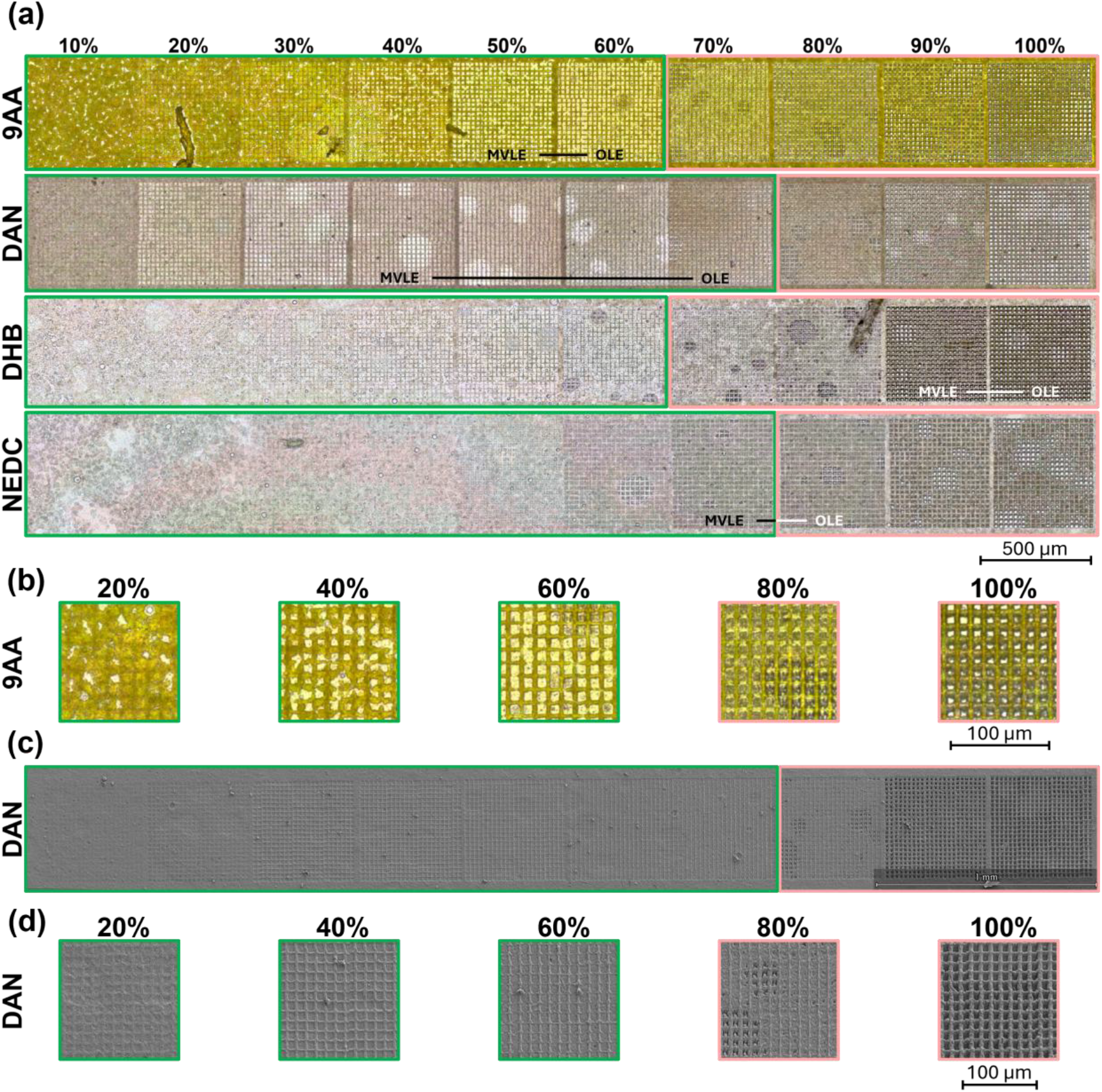
(a) Tiled microscope images of ablation areas for each tested matrix, using 20 µm pixel sizes and no post-ionisation, across all laser energies, with overlays indicating the MVLE–OLE range for each matrix as previously determined spectrally. (b) Zoom-ins of regions in the 9AA optical images for a selection of laser energy settings. (c) Tiled SEM micrographs of ablation areas in sample coated with DAN. (d) Zoom-ins of regions in the SEM micrographs of the sample coated with DAN for a selection of laser energy settings. For all images, the green box outlines the microscope images of analysis areas where the damage to the PEN membrane underneath the matrix and sample appears minimal, while the other ablation area images have a red box around them.

Since the MALDI MS instrument laser was set up to raster along three parallel lines inside each 20 µm pixel (a.k.a “beam scan”), the microscope images were assessed for intra-pixel laser raster lines, accompanied by a change in colour (*e*.*g*. for 9AA the change is from yellow to white between 20 and 60%) which suggested there was no more matrix. Typically, the colour change comes first, followed by the appearance of darker patches which, as confirmed by the SEM, are due to ablation of the surface of the polymer substrate. The images then tend to get brighter as the membrane becomes more fully removed, with near-total removal at 100% laser energy. For the 5 µm pixels, the images were more difficult to assess but evidence of colour change in the centre of the pixel was still visible, with the same pattern as for the samples acquired at 20 µm pixel size. MVLE and OLE ranges determined previously from ion data are overlayed on the microscope images. For both 9AA and DAN, the full MVLE to OLE range is within the damage limit. For NEDC, only the MVLE was within, and for DHB no laser energies were within these bounds. This makes DHB unsuitable for use with these substrates within a multimodal workflow since even the MVLE % energy causes significant substrate damage.

The potential protective effect to the substrate of the matrices which have greater laser fluence thresholds (NEDC and DHB) was also considered. Although the 5 µm MALDI images (SI Figure S6) could support this, when all ablation areas across all samples are considered, there is no correlation. It is possible that using matrix application techniques which produce more even matrix layers, such as sublimation, or depositing a thicker layer of matrix may be better at reducing laser energy reaching the substrate. For the datasets without post-ionisation, all the damage limits were either 60 or 70% laser energy, regardless of pixel size. The limit for DAN increased by 10% at the higher pixel size, while the limit for DHB reduced by 10% and no change was noted for NEDC and 9AA. In the MALDI-2 datasets, the damage limits were either 30 or 40% at 5 µm pixel size and all 30% at 20 µm pixel size. This is slightly lower than the limits without post-ionisation, even when adjusting for the automatic 15% laser energy boost. This difference may be attributed to the lower laser frequency (repetition rate) used in the MALDI-2 experiments (SI Table S1), since the number of laser shots was kept constant for both ionisation modes. Operating the laser at a lower repetition rate of 1 kHz could increase the energy per pulse and therefore transfer more total energy to the sample^30, 31^.

SEM imaging (Figure 4, SI Figure S6) was used to add further context to the observations of damage via assessment of optical images for DAN-coated samples. Extensive tissue removal and possible membrane damage is clearer in SEM micrographs because the higher resolution allows the laser craters to become visible in detail. The first signs of membrane damage are seen where there are pre-existing holes in the tissue, which look like faint circular and oval shapes that become increasingly bright in the 10 to 70% laser energy regions of Figure 4. They become more visible in the 80 to 100% laser energy regions due to the three dark, vertical raster lines in each pixel, consistent with how the instrument ‘beam scan’ setting works. The SEM micrographs supported the damage limits assigned from the optical microscopy.

Based on the combined assessment of the relative ions of interest intensities, spectral quality and MVLE to OLE range within the context of the sample damage limit, DAN is selected as the most appropriate matrix. It also has the advantage of being a dual-polarity MALDI matrix^32^ which has been used to image lipids and peptides^33^ as well as small molecules^34^, making it a versatile choice. One of its drawbacks is that it is more hazardous than most matrices, which requires careful handling with additional health and safety measures.

### Substrate comparison

The optimisation experiments described above were conducted on samples with gPEN as the substrate, a commercially available glass-supported PEN membrane substrate. Another format, fPEN, consists of PEN membrane supported by a metal frame. These substrates offer compatibility for different modalities; for example, gPEN is not directly suitable for mounting samples for X-ray fluorescence (XRF) or particle-induced x-ray emission (PIXE) analysis due to the glass support, whereas fPEN is suitable.

The recommended substrate for the mass spectrometer is a conductive ITO slide so using non-conductive membrane substrates was expected to produce lower intensities. As a compromise, the membrane from an fPEN slide was glued then cut out onto an ITO slide (itoPEN; see Materials & Methods section and SI Figure S3), which could allow better charge dissipation than gPEN.

The substrates gPEN, fPEN, itoPEN and ITO were compared using the same combinations of pixel sizes and ionisation modes as for the experiments discussed above. Only the range of laser energies between the MVLE and OLE (as previously defined on gPEN) for each set of parameters was acquired. Figure 5 compares the substrates with the relative intensities of the same six ion of interest as used in the previous analysis.

**Figure 5.**
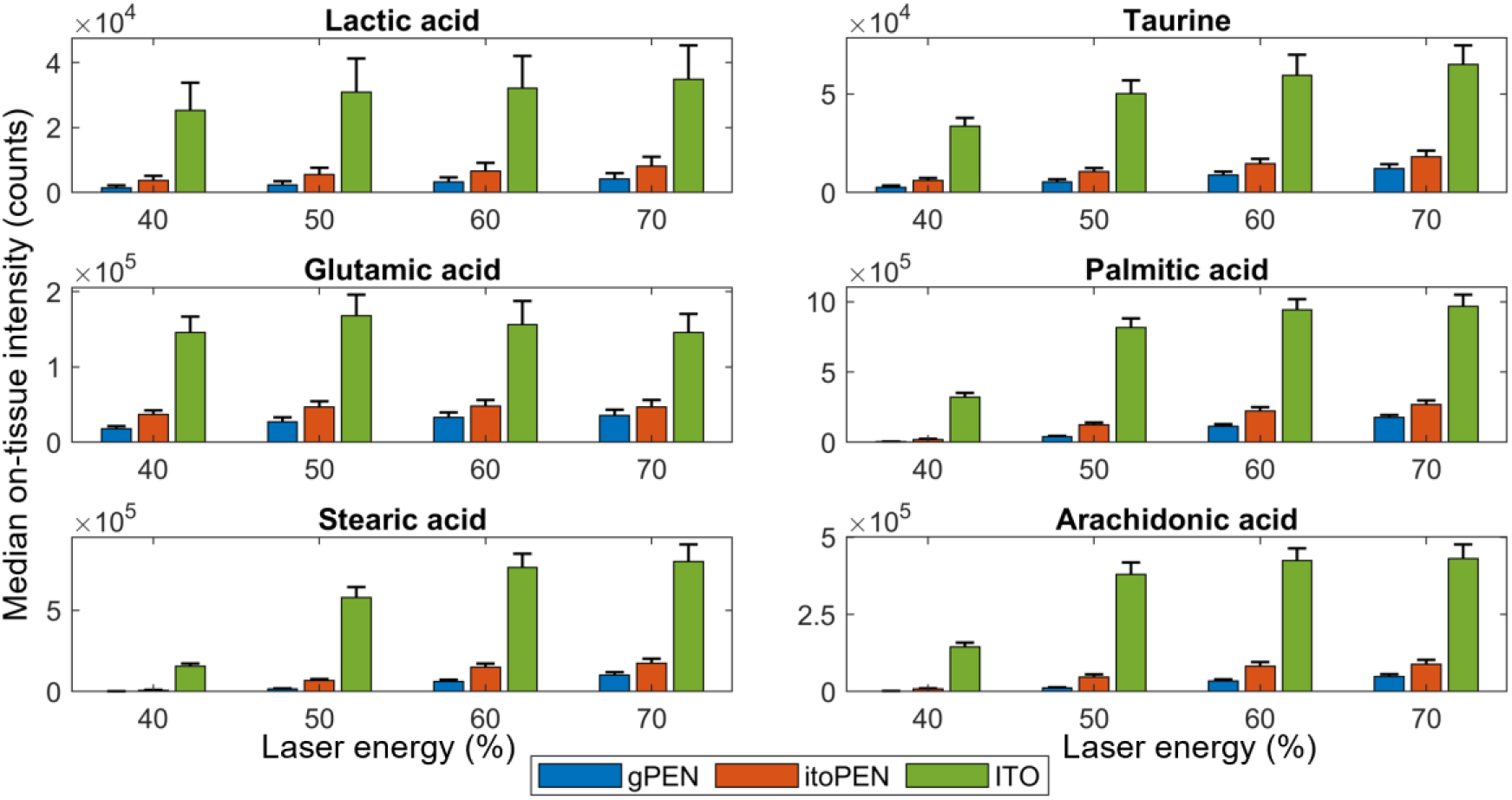
Grouped bar graphs comparing the median on-tissue intensities of the deprotonated peaks of six analytes of interest for each substrate and all MALDI 20 µm datasets.

No signal above the noise level was detected from fPEN at any tested laser energy; these datasets were therefore excluded from subsequent analysis and are not shown in Figure 5. The weak ablation marks on fPEN suggested that the sample surface was beyond the effective focal range of the laser. This is likely attributable to way the fPEN sample was loaded into the instrument, which required fixing it to the surface of a MALDI spot plate with copper tape (see Materials & Methods section), since neither the substrate nor the slide holder could be resized. It is possible there was incomplete contact between the membrane and the MALDI target plate, due to the metal frame being out of alignment with the horizontal plane. This would position the sample further above the surface of the steel MALDI target plate, potentially compounded by membrane thickness (4 µm). Although the stage permits a degree of vertical adjustment via built in height calibration, the focal position is highly sensitive to height variation because the laser spot diameter is approximately 4 µm in FWHM. MALDI-2 acquisition was also prevented by obstruction of the MALDI-2 laser path by the metal frame.

The remaining substrates generated sufficient signal for comparison. Without post-ionisation (Figure 5), signal intensity was highest on ITO, followed by itoPEN and then gPEN. This trend is consistent with the expected effect of substrate conductivity whereby reduced conductivity promotes surface charge accumulation and reduces ion generation and extraction efficiency^35-37^. Although mounting a PEN membrane on ITO does not alter the intrinsic conductivity of the membrane itself, the conductive support may facilitate charge dissipation.

With MALDI-2 enabled, substrate-dependent trends were less consistent. The ion intensities from each substrate that were more similar, particularly at the 5 µm pixel size (SI Figure S7). Between 10 and 30% laser energy, gPEN produced the highest intensities for more ions than the other substrates, whereas above 30%, ITO produced the highest intensities overall (followed by gPEN, then itoPEN). MALDI-2 seems to recover ion signal from the PEN substrate relative to ITO suggesting that despite insulating substrates quenching of ion signal in MALDI, there remains sufficient gas phase material conducive to MALDI-2 post-ionisation to produce viable or even high-quality spectral information from PEN in various forms. The poor thermal conduction of UV-absorbing PEN^21, 38^ may promote localised heating during laser irradiation, increasing desorption^39^ and producing a larger population of gas-phase neutrals available for MALDI-2 post-ionisation^40, 41^.

To assess both the overall spectral quality and the extent of any substrate-derived background peak contributions, SI Figure S8 shows the on-tissue spectra for the same datasets discussed above at the MVLE and OLE. At the OLE, MALDI without post-ionisation produced comparable spectral profiles across ITO, itoPEN and gPEN, with the ions of interest readily detected on all three substrates. However, absolute signal intensities were consistently highest on ITO, followed by itoPEN and gPEN. With MALDI-2 enabled, several additional peaks were observed in the OLE spectra from both PEN-based substrates, particularly above *m/z* 400. The predominantly off-tissue localisation of these features was consistent with a membrane-derived origin, although they did not substantially alter the overall spectral profiles or interfere with detection of the ions of interest. They do, however, highlight the importance of defining an upper laser energy threshold for analyses on PEN. Overall, these results indicate that both PEN-based substrates can generate good-quality spectra that are broadly comparable to those acquired on ITO.

Selection between gPEN and itoPEN for a multimodal study will depend on a number of factors, including the modalities intended to be included, MALDI laser setting, ionisation mode and time available for sample preparation. Without MALDI-2, itoPEN produces higher intensities, at the cost of longer sample preparation time due to the membrane transfer step. In MALDI-2 mode, at low laser energy settings, the gPEN slides may produce higher intensities, whereas at higher laser energies, itoPEN performs better – although it should be noted that both differences are marginal. The membrane transfer step may also introduce additional variability through membrane stretching and incomplete flattening against the ITO. Due to these factors, gPEN was selected as the substrate for mouse brain imaging, with ITO included for reference.

### Mouse brain and cell imaging

Serial coronal sections were acquired at the MVLE and OLE using MALDI and MALDI-2 at 20 µm pixel size. Figure 6 shows single ion images for a range of putatively assigned, deprotonated analyte peaks.

**Figure 6.**
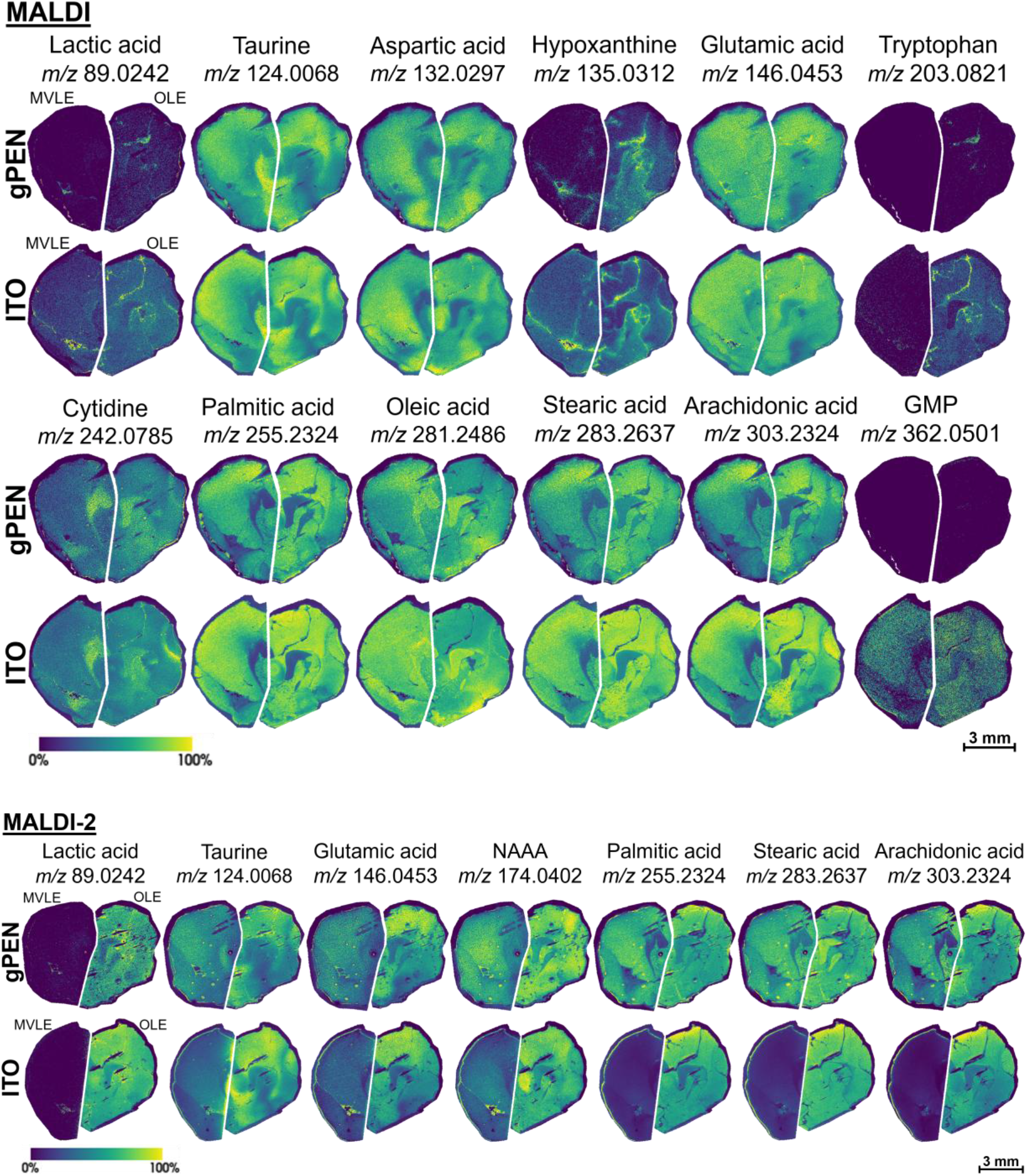
Single ion images of coronal mouse brain mounted on gPEN and ITO. Assignments are putative and the images were created with a ±15 ppm mass error around the deprotonated ion mass. Approximately half of each brain was acquired using the MVLE for that pixel size, and the other half using the OLE. For MALDI datasets, the OLE is 70% and MVLE is 40%, and for MALDI-2 they are 70% and 10% respectively. Each half-brain is on an independent colour scale, with the upper intensity limit thresholded to the 99% quantile of all image intensities for clarity. GMP = guanosine monophosphate and NAA = N-acetylaspartic acid.

**Figure 7.**
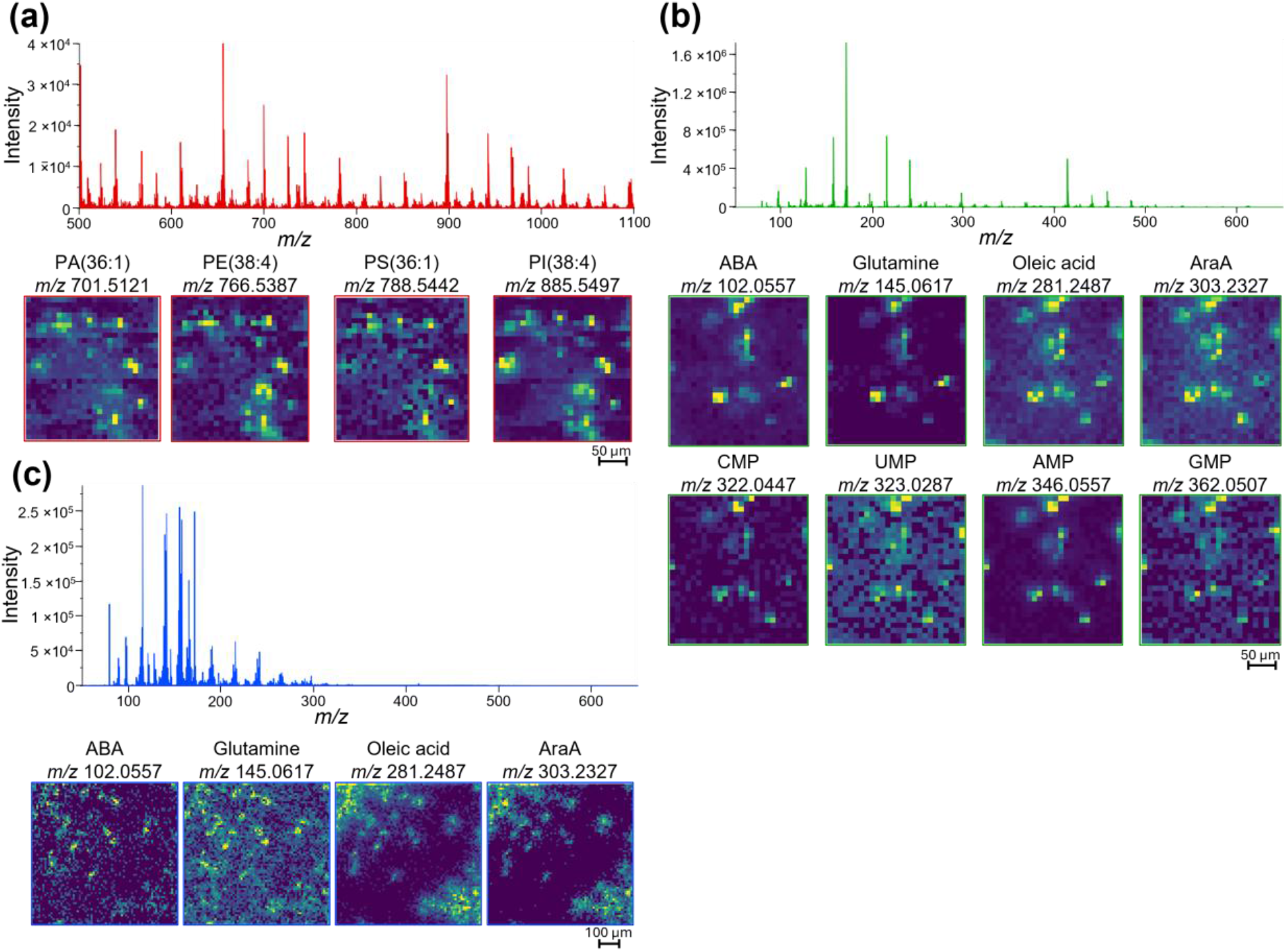
Mean spectra from pixels on PANC-1 cells acquired with MALDI-2 and 10 µm pixel size and accompanying single ion images. (a) Lipid region data from cells grown on fPEN and acquired on itoPEN. (b) Metabolite region data from cells grown on fPEN and acquired on itoPEN. (c) Metabolite region data from cells grown and acquired on gPEN. Assignments are putative and the images were created with a ±15 ppm mass error around the deprotonated ion mass. The upper intensity limit of all images was thresholded to the 99% quantile of all pixel intensities for clarity. PA = phosphatidic acid, PE = phosphatidylethanolamine, PS = phosphatidylserine, PI = phosphatidylinositol, ABA = aminobutyric acid, AraA = arachidonic acid, CMP = cytidine monophosphate, UMP = uridine monophosphate, AMP = adenosine monophosphate, GMP = guanosine monophosphate.

Peaks were putatively assigned to analytes across a range of molecular classes, including amino acids, nucleosides, carbohydrates and carboxylic acids. Despite the reduced intensities at the MVLE, many matched peaks retained spatial distributions comparable to those observed at the OLE. As expected, more spatially resolved ion images were obtained at the OLE. Without post-ionisation, gPEN and ITO produced comparable image quality for ions detected in both, although the absolute intensities and number of detected ions were generally higher on ITO.

In MALDI-2 mode, the ITO MVLE region showed poorer image quality than the corresponding gPEN region. The edge of the tissue is much brighter than the rest of the tissue and there is reduced visibility of internal brain features. The remaining acquisition conditions produced comparable images and retained tissue features. Substrate-dependent intensity differences varied between ions, although most ions showed higher signal on ITO. Overall, these results show that gPEN can preserve spatially resolved molecular information in mouse brain tissue, despite lower absolute signal intensity than ITO.

Applications requiring the analysis of individual microorganisms or animal cells depend on sufficiently high spatial resolution to distinguish neighbouring cells^42, 43^. This is difficult to achieve with MALDI on any substrate, but perhaps more so on non-conductive slides since the sensitivity is lower and will be compounded by sampling less material per pixel. Since it was shown that using MALDI-2 is minimally affected by the choice of substrate, this was used to analyse PANC-1 cells grown on gPEN and fPEN (later made into itoPEN prior to MALDI analysis).

The mean spectra created from the cell regions have many peaks that are similarly intense to those acquired on tissue with bigger pixel sizes (Figure 3). The single ion images show that many of these peaks also produce single ion images with defined cell outlines. Two different cell fixation methods were used, which had a strong influence on spectral distributions. Using PFA resulted in lower overall intensities (∼10-fold) as well as a concentration of the signal to between

*m/z* 50 and 350, while using freeze drying produced spectra with signal distributed throughout the entire *m/z* 50 to 650 range. This allowed analytes such as ribonucleotides to be detected (CMP, UMP, AMP, GMP). It should be noted that the PFA-fixed cells were analysed on gPEN while the freeze-dried cells were analysed on itoPEN, which partially contributes to the difference in intensities, as discussed previously. It was also shown to be possible to image lipid distributions at single-cell resolution using the same sample preparation.

## Conclusions

This study demonstrates how MALDI MSI can be used on samples with PEN membranes as the substrate, to enable multimodal analysis on the same tissue section. By evaluating matrix choice, laser energy, pixel size, ionisation mode and substrate format, we demonstrated that optimisation on PEN must balance ion intensity and image quality with sample and membrane damage. The use of MVLE and OLE provided a practical framework for assessing acquisition parameters.

Among the matrices tested, DAN gave the strongest overall performance, with higher ion intensities, lower MVLE values and a wider viable laser energy range. Although other matrices performed better in specific analyte cases, DAN presented the best compromise. It also happens to be a versatile matrix, which other publications have shown to work in both polarities and for a wide range of molecules.

Substrate selection was also shown to have substantial influence on MALDI performance. Under the tested configuration, fPEN was not suitable for MALDI analysis on the TimsTOF flex instrument. gPEN and itoPEN both produced lower intensities than ITO, particularly without MALDI-2 post-ionisation, but offered different trade-offs. itoPEN may be preferable when MALDI is the final modality and higher ion intensity is prioritised, whereas gPEN may be preferable when lower laser energies, simpler preparation or reduced transfer-related variability are more important. Substrate choice may also be constrained by compatibility with downstream modalities and therefore influence the order of analysis. Application of the optimised method on gPEN to mouse brain tissue confirmed that imaging on PEN substrates produces comparable ion images to ITO. Furthermore, it was shown to be possible to image the metabolites and lipids of single PANC-1 cells, using different cell fixation methods on both gPEN and itoPEN substrates.

Overall, these results provide guidance for matrix, instrument parameter and substrate selection for same-section multimodal analysis pipelines using MALDI analysis on PEN membranes.

