## Supporting Information for "Optimising MALDI on PEN substrates for small molecule analysis in the context of same-section multimodal workflows"

Table S1 – Instrument parameters for all methods

| Parameter | MALDI 5 µm | MALDI 20 µm | MALDI-2 5 µm | MALDI-2 20 µm |
|---|---|---|---|---|
| Mass range | 50-650 | 50-650 | 50-650 | 50-650 |
| Polarity | Negative | Negative | Negative | Negative |
| Postionisation | Off | Off | On | On |
| Trigger Delay / µs | N/A | N/A | 5.0 | 5.0 |
| Shots | 15 | 500 | 15 | 500 |
| Laser frequency | 10000 | 10000 | 1000 | 1000 |
| MALDI plate offset / V | 50.0 | 50.0 | 50.0 | 50.0 |
| Deflection delta / V | -45.0 | -45.0 | -45.0 | -45.0 |
| Funnel 1 RF / Vpp | 125.0 | 125.0 | 125.0 | 125.0 |
| isCID energy / eV | -0.0 | -0.0 | -0.0 | -0.0 |
| Funnel 2 RF / Vpp | 100.0 | 100.0 | 100.0 | 100.0 |
| Multipole RF / Vpp | 200.0 | 200.0 | 200.0 | 200.0 |
| Collision energy / eV | 8.0 | 8.0 | 8.0 | 8.0 |
| Collision RF / Vpp | 500.0 | 500.0 | 500.0 | 500.0 |
| Ion energy / eV | 5.0 | 5.0 | 5.0 | 5.0 |
| Low mass / m/z | 50.0 | 50.0 | 50.0 | 50.0 |
| Transfer time / µs | 40.0 | 40.0 | 40.0 | 40.0 |
| Pre-pulse storage / µs | 5.0 | 5.0 | 5.0 | 5.0 |

Table S2 – List of ions of interest with accompanying information

| Name | Monoisotopic mass | Chemical formula | Adduct | Theoretical *m/z* adduct |
|---|---|---|---|---|
| Lactic acid | 90.03169 | C3H6O3 | [M-H] | 89.02387 |
| Taurine | 125.0147 | C2H7NO3S | [M-H] | 124.0077 |
| Glutamic acid | 147.0532 | C5H9NO4 | [M-H] | 146.0453 |
| Palmitic acid | 256.2402 | C16H32O2 | [M-H] | 255.2324 |
| Stearic acid | 284.2715 | C18H36O2 | [M-H] | 283.2637 |
| Arachidonic acid | 304.2402 | C20H32O2 | [M-H] | 303.2324 |

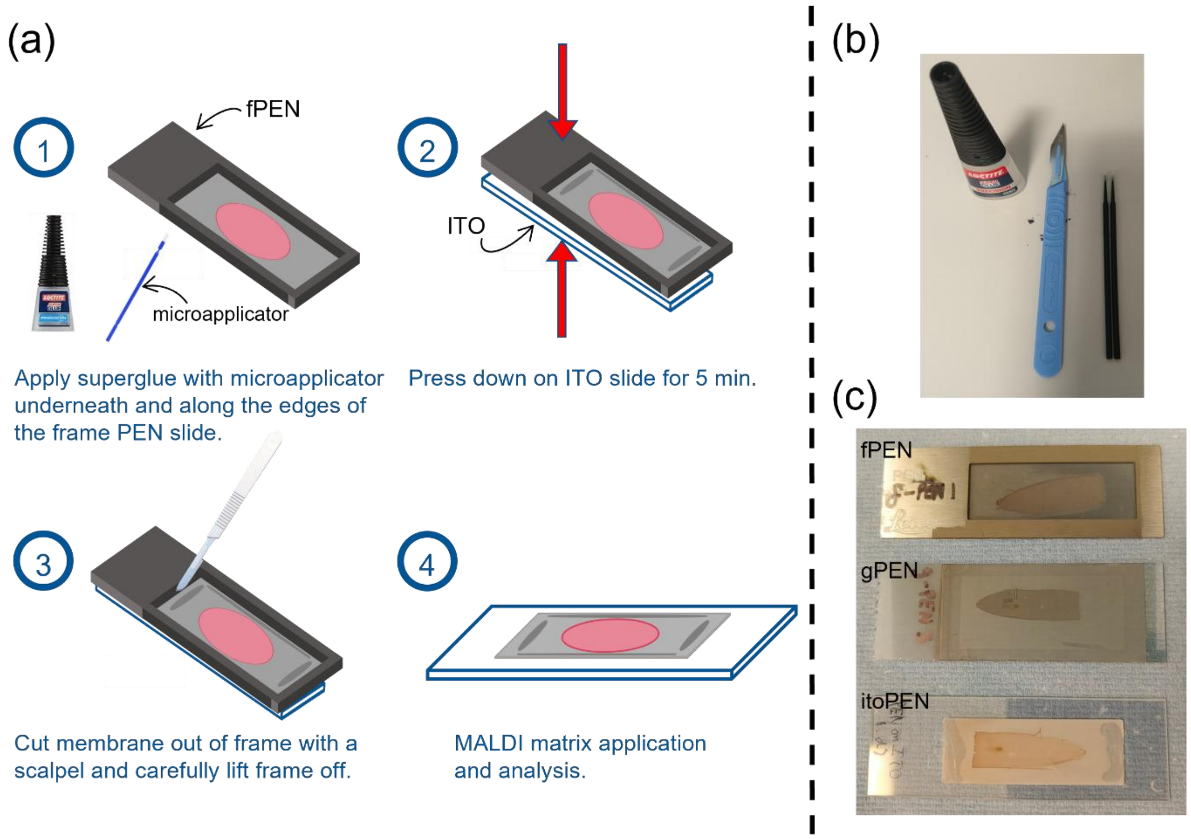


Figure S3 – (a) Membrane transfer procedure. (b) Equipment required for membrane transfer procedure. (c) Picture of all PEN slides prior to MALDI analysis.

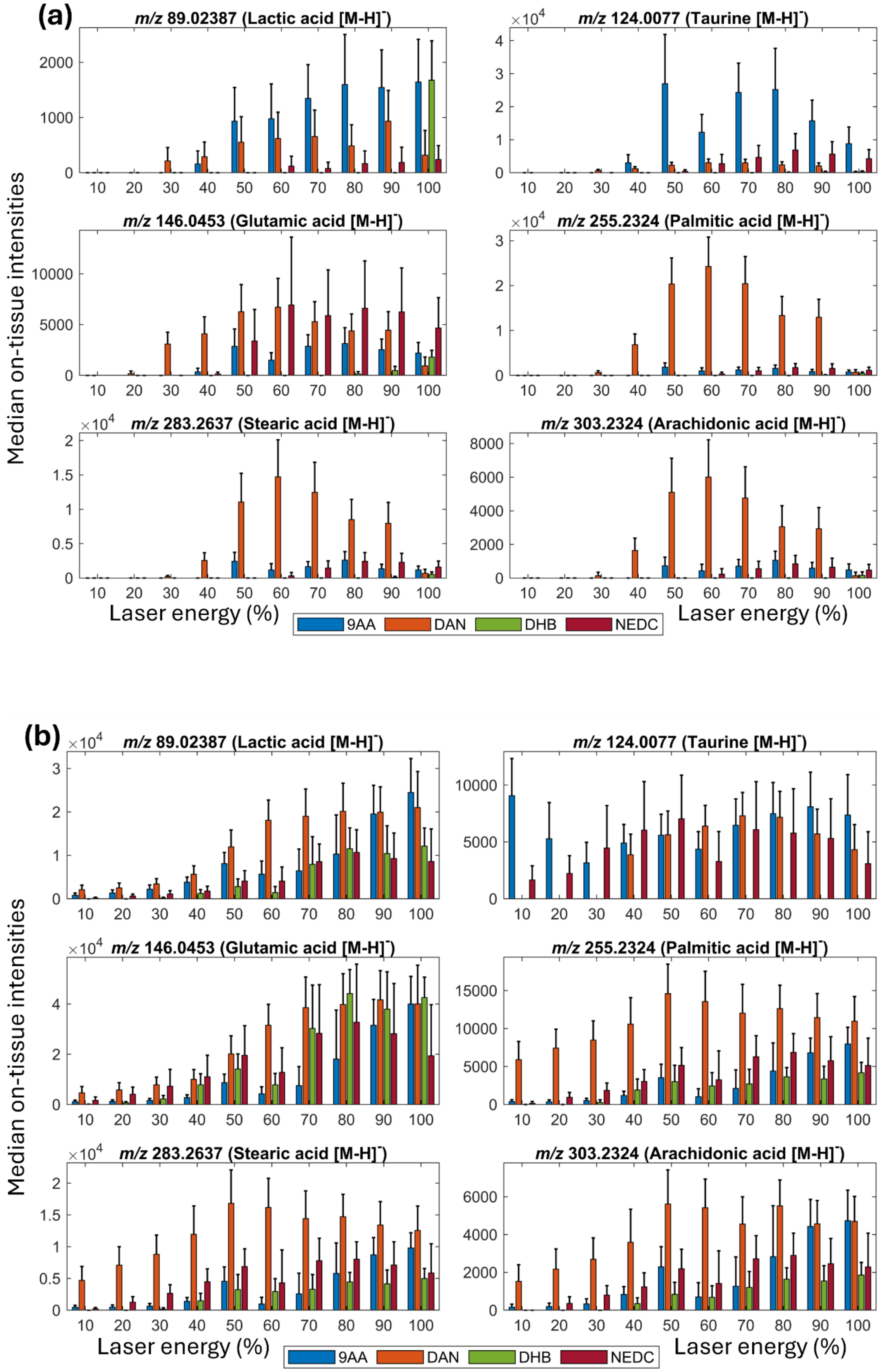
(a)
m/z 89.02387 (Lactic acid [M-H]-)
m/z 124.0077 (Taurine [M-H]-)
m/z 146.0453 (Glutamic acid [M-H]-)
m/z 255.2324 (Palmitic acid [M-H]-)
m/z 283.2637 (Stearic acid [M-H]-)
m/z 303.2324 (Arachidonic acid [M-H]-)
Median on-tissue intensities
Laser energy (%)
9AA
DAN
DHB
NEDC
(b)
m/z 89.02387 (Lactic acid [M-H]-)
m/z 124.0077 (Taurine [M-H]-)
m/z 146.0453 (Glutamic acid [M-H]-)
m/z 255.2324 (Palmitic acid [M-H]-)
m/z 283.2637 (Stearic acid [M-H]-)
m/z 303.2324 (Arachidonic acid [M-H]-)
Median on-tissue intensities
Laser energy (%)
9AA
DAN
DHB
NEDC

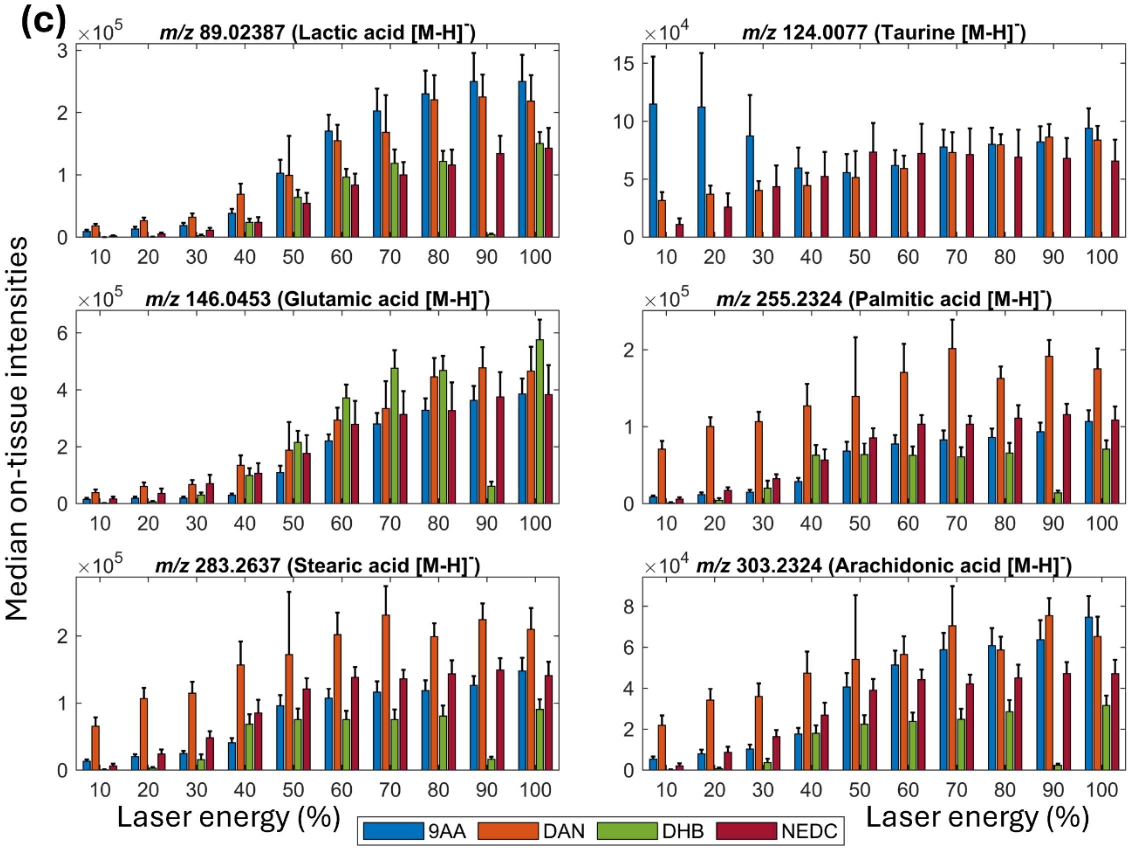


Figure S4 – Grouped bar charts of median on-tissue intensities across the working range of instrument laser energy settings for various deprotonated ions. Different coloured bars for each matrix are shown at each laser energy. (a) MALDI, 5 µm pixel data (without postionisation) on gPEN. (b) MALDI-2, 5 µm pixel data. (c) MALDI-2, 20 µm pixel data

Table S5 – Table of polyethylene naphthalate fragment peaks and whether a match was found in the MVLE and OLE spectra of DAN-coated samples. Matching performed using 15 ppm tolerance.

| Structure | * 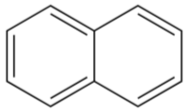 | 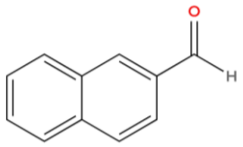 | 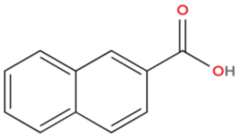 |
|---|---|---|---|
| Formula | $C_{10}H_8$ | $C_{11}H_8O$ | $C_{11}H_8O_2$ |
| Adduct | [M-H]- | [M-H]- | [M-H]- |
| Expected *m/z* | 127.0553 | 155.0502 | 171.0451 |
| MALDI, 5µm, OLE (80%), 9AA measured *m/z* | 127.0548 (-3.9 ppm) | 155.0492 (-6.4 ppm) | 171.0449 (-1.2 ppm) |
| MALDI, 20µm, OLE (60%), 9AA measured *m/z* | Not detected | Not detected | 171.0446 (-2.9 ppm) |

| | | | |
|---|---|---|---|
| MALDI, 5µm, OLE (60%), DAN measured *m/z* | Not detected | Not detected | 171.0442 (-5.3 ppm) |
| MALDI, 20µm, OLE (70%), DAN measured *m/z* | Not detected | Not detected | 171.0446 (-2.9 ppm) |
| MALDI, 5µm, OLE (100%), DHB measured *m/z* | Not detected | 155.0498 (-2.6 ppm) | 171.0447 (-2.3 ppm) |
| MALDI, 20µm, OLE (100%), DHB measured *m/z* | Not detected | 155.0498 (-2.6 ppm) | 171.0447 (-2.3 ppm) |
| MALDI, 5µm, OLE (80%), NEDC measured *m/z* | Not detected | 155.0482 (-12.9 ppm) | 171.0444 (-4.3 ppm) |
| MALDI, 20µm, OLE (80%), NEDC measured *m/z* | 127.0546 (-2.4 pm) | 155.0496 (-3.9 ppm) | 171.0446 (-2.9 ppm) |
| MALDI-2, 5µm, OLE (70%), 9AA measured *m/z* | 127.0558 (+7.1 ppm) | 155.0504 (+1.3 ppm) | 171.0453 (+1.2 ppm) |
| MALDI-2, 20µm, OLE (70%), 9AA measured *m/z* | 127.0557 (+6.3 ppm) | 155.0506 (+2.6 ppm) | 171.0456 (+2.9 ppm) |
| MALDI-2, 5µm, OLE (60%), DAN measured *m/z* | 127.0552 (+2.4 ppm) | 155.0505 (+1.9 ppm) | 171.0446 (-2.9 ppm) |
| MALDI-2, 20µm, OLE (70%), DAN measured *m/z* | 127.0554 (3.9 ppm) | 155.0511 (+5.8 ppm) | 171.0455 (2.3 ppm) |
| MALDI-2, 5µm, OLE (70%), DHB measured *m/z* | 127.0554 (3.9 ppm) | 155.0500 (-1.3 ppm) | 171.0449 (-1.2 ppm) |
| MALDI-2, 20µm, OLE (70%), DHB measured *m/z* | 127.0557 (+6.3 ppm) | 155.0507 (+3.2 ppm) | 171.0447 (-2.3 ppm) |
| MALDI-2, 5µm, OLE (70%), NEDC measured *m/z* | 127.0553 (+3.1 ppm) | 155.0498 (-2.6 ppm) | 171.0447 (-2.3 ppm) |
| MALDI-2, 20µm, OLE (60%), NEDC measured *m/z* | 127.0554 (3.9 ppm) | 155.0505 (+1.9 ppm) | 171.0455 (2.3 ppm) |

* This fragment could originate from 1,5-diaminonaphthalene (DAN) or *N*-(1-Naphthyl)ethylenediamine dihydrochloride (NEDC) matrices

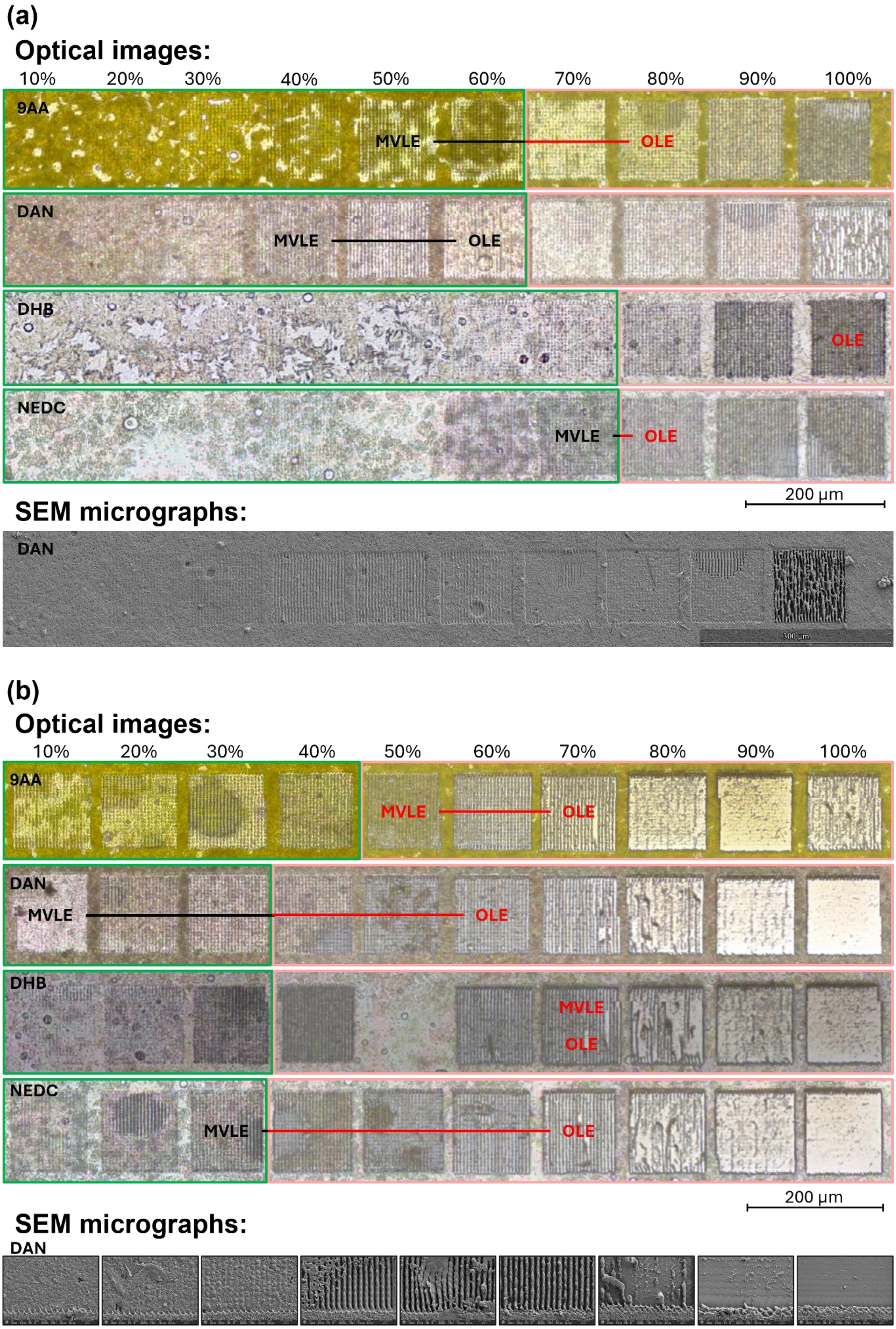
(a)
Optical images:
10%
20%
30%
40%
50%
60%
70%
80%
90%
100%
9AA
MVLE
OLE
DAN
MVLE
OLE
DHB
OLE
NEDC
MVLE
OLE
200 µm
SEM micrographs:
DAN
(b)
Optical images:
10%
20%
30%
40%
50%
60%
70%
80%
90%
100%
9AA
MVLE
OLE
DAN
MVLE
OLE
DHB
MVLE
OLE
NEDC
MVLE
OLE
200 µm
SEM micrographs:
DAN

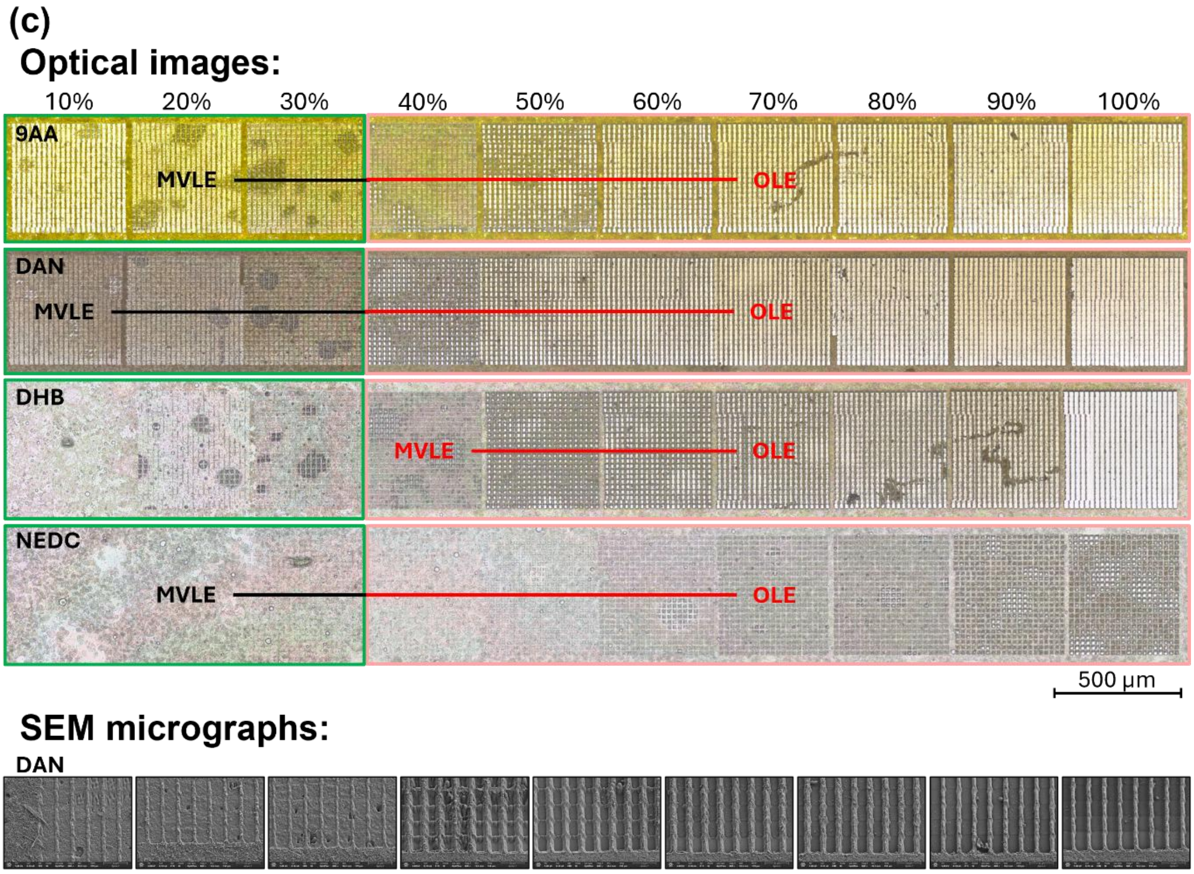


Figure S6 - (a) Top: Tiled microscope images of ablation areas for each tested matrix, using 5 µm pixel sizes and no postionisation, across all laser energies, with overlays indicating the MVLE–OLE range for each matrix. Bottom: Tiled SEM micrographs of the same ablation areas in DAN matrix. (b) Top: Tiled microscope images of ablation areas for each tested matrix, using 5 µm pixel sizes and MALDI-2 postionisation, across all laser energies, with overlays indicating the MVLE–OLE range for each matrix. Bottom: SEM micrographs of the same ablation areas in DAN matrix. (c) Top: Tiled microscope images of ablation areas for each tested matrix, using 20 µm pixel sizes and MALDI-2 postionisation, across all laser energies, with overlays indicating the MVLE–OLE range for each matrix. Bottom: Tiled SEM micrographs of the same ablation areas in DAN matrix.

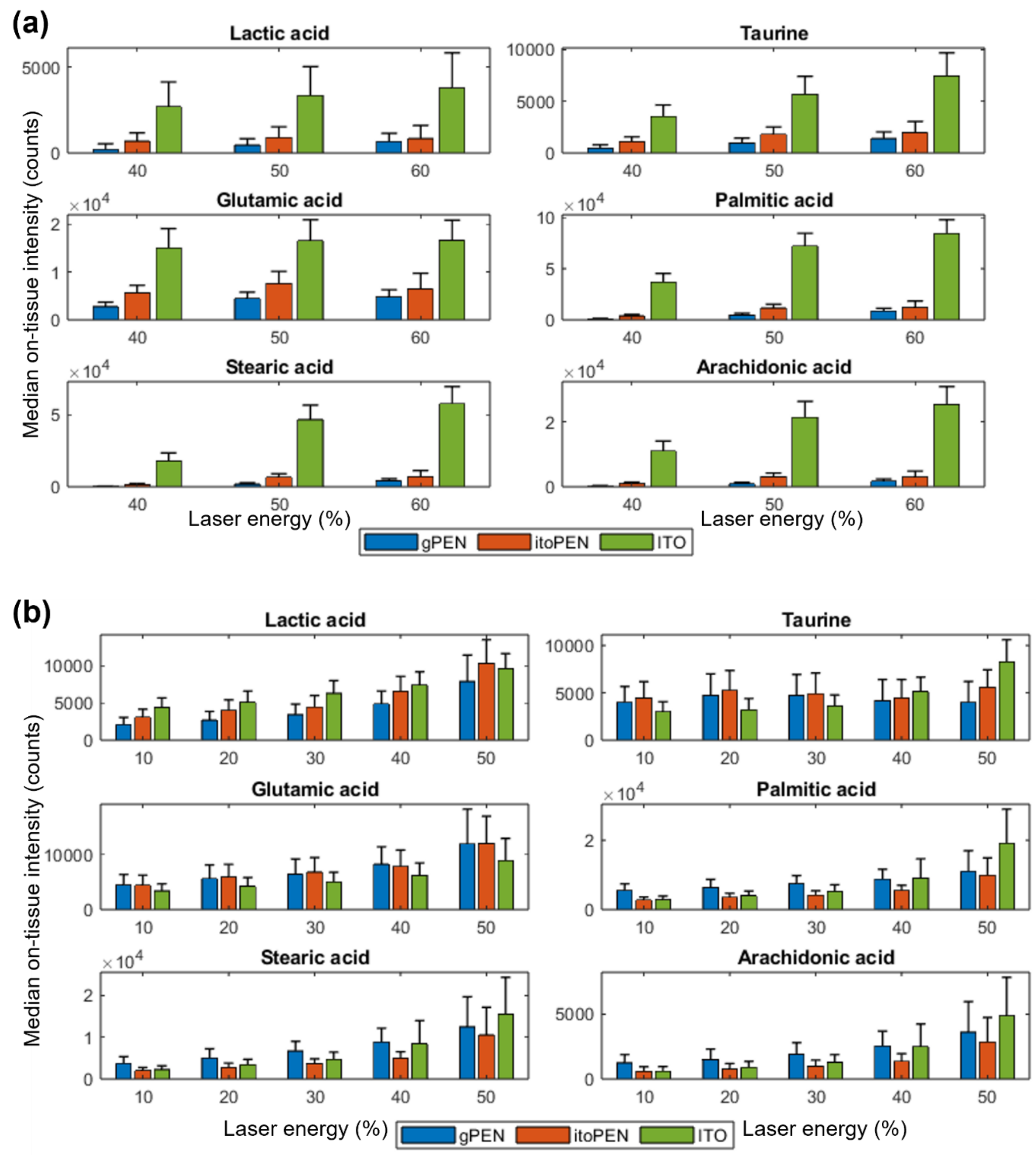
(a)
Lactic acid
Taurine
Glutamic acid
Palmitic acid
Stearic acid
Arachidonic acid
Median on-tissue intensity (counts)
Laser energy (%)
gPEN
itoPEN
ITO
(b)
Lactic acid
Taurine
Glutamic acid
Palmitic acid
Stearic acid
Arachidonic acid
Median on-tissue intensity (counts)
Laser energy (%)
gPEN
itoPEN
ITO

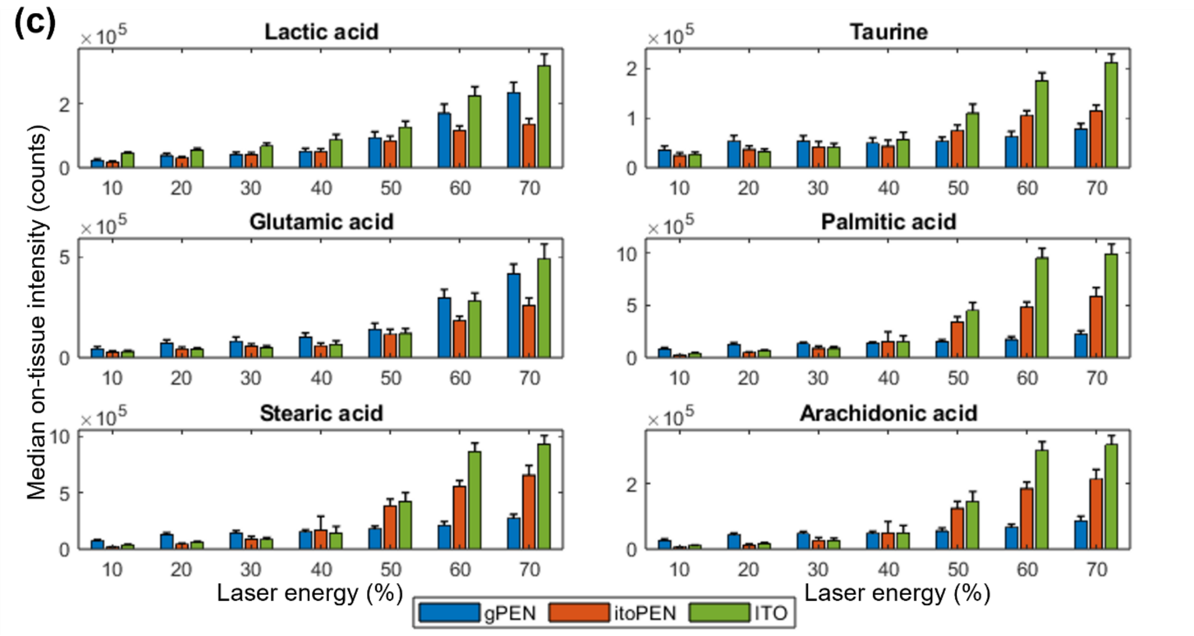


Figure S7– Grouped bar graphs comparing the median on-tissue intensities of the deprotonated peaks of six analytes of interest for each substrate and (a) all MALDI 5 µm datasets, (b) all MALDI-2 5 µm datasets, (c) all MALDI-2 20 µm datasets.

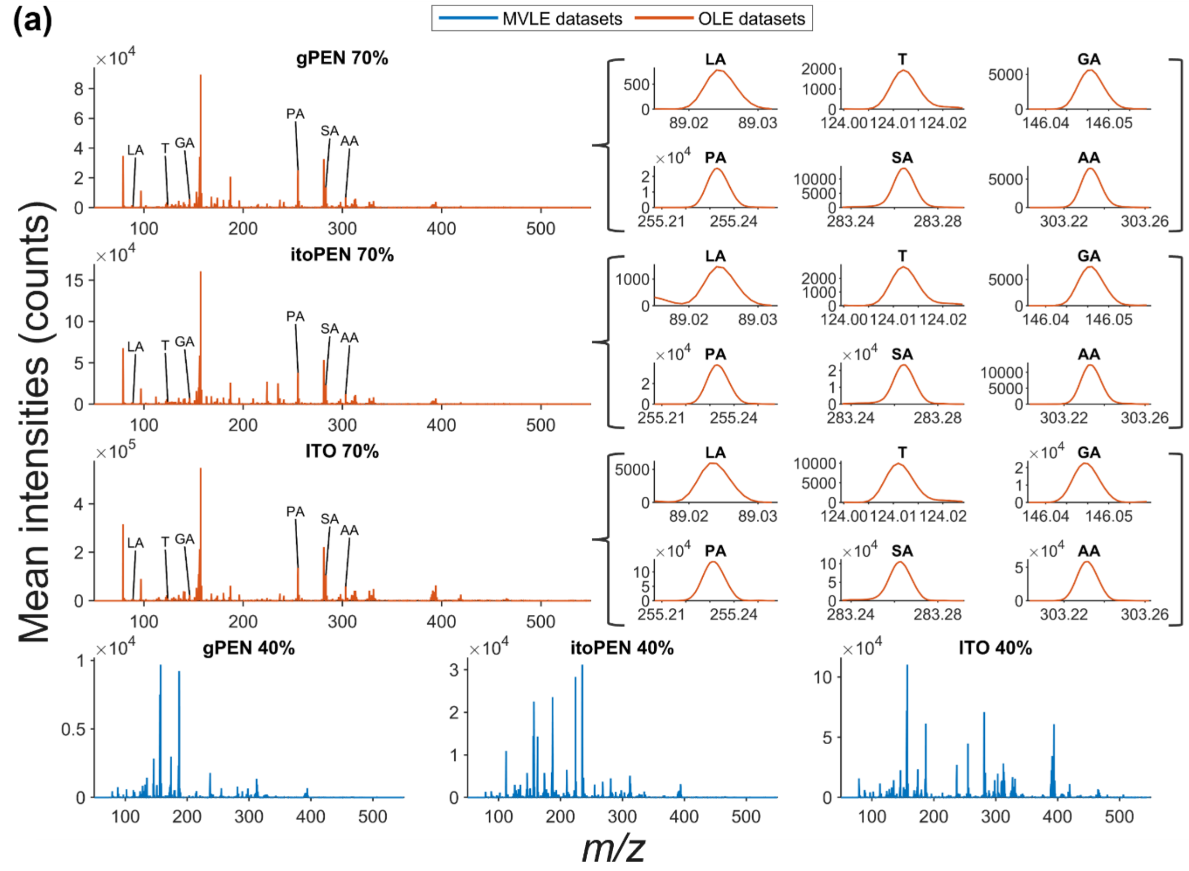

(b)

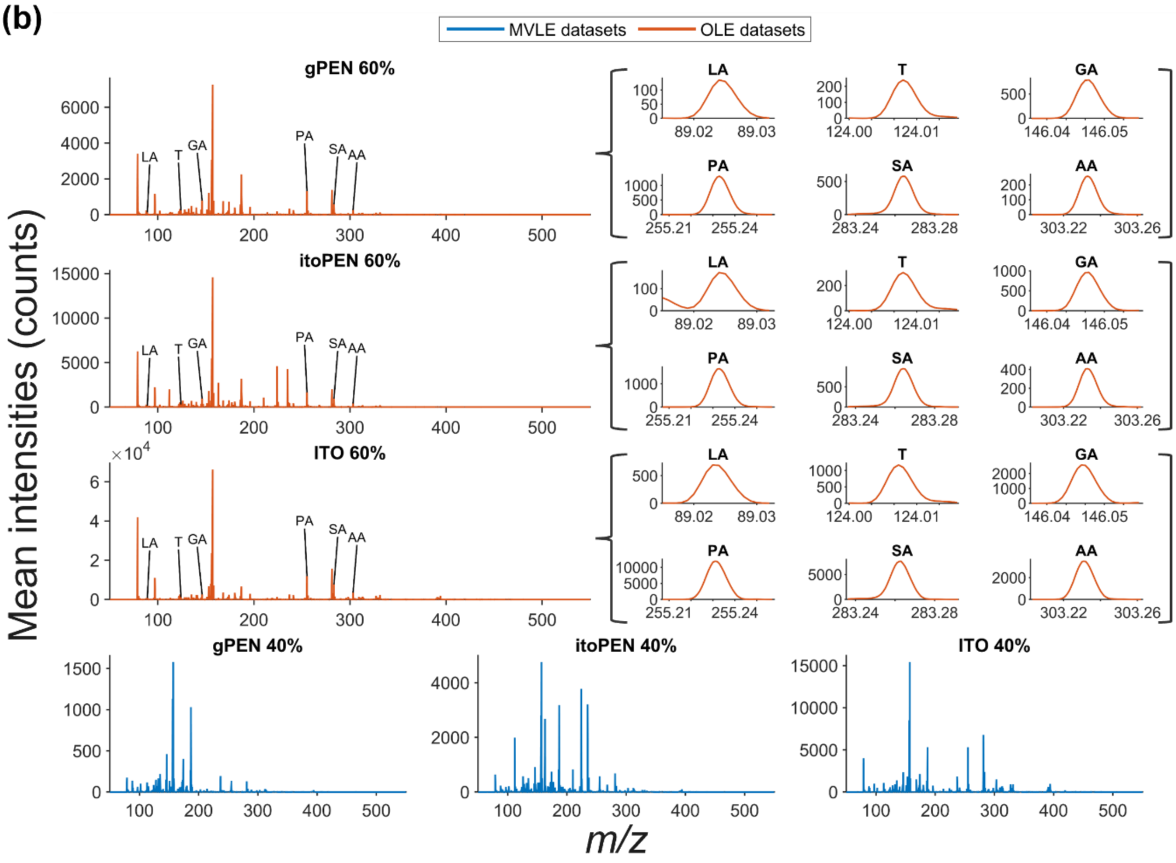
MVLE datasets
OLE datasets
gPEN 60%
itoPEN 60%
ITO 60%
gPEN 40%
itoPEN 40%
ITO 40%
LA
T
GA
PA
SA
AA
Mean intensities (counts)
m/z

(c)

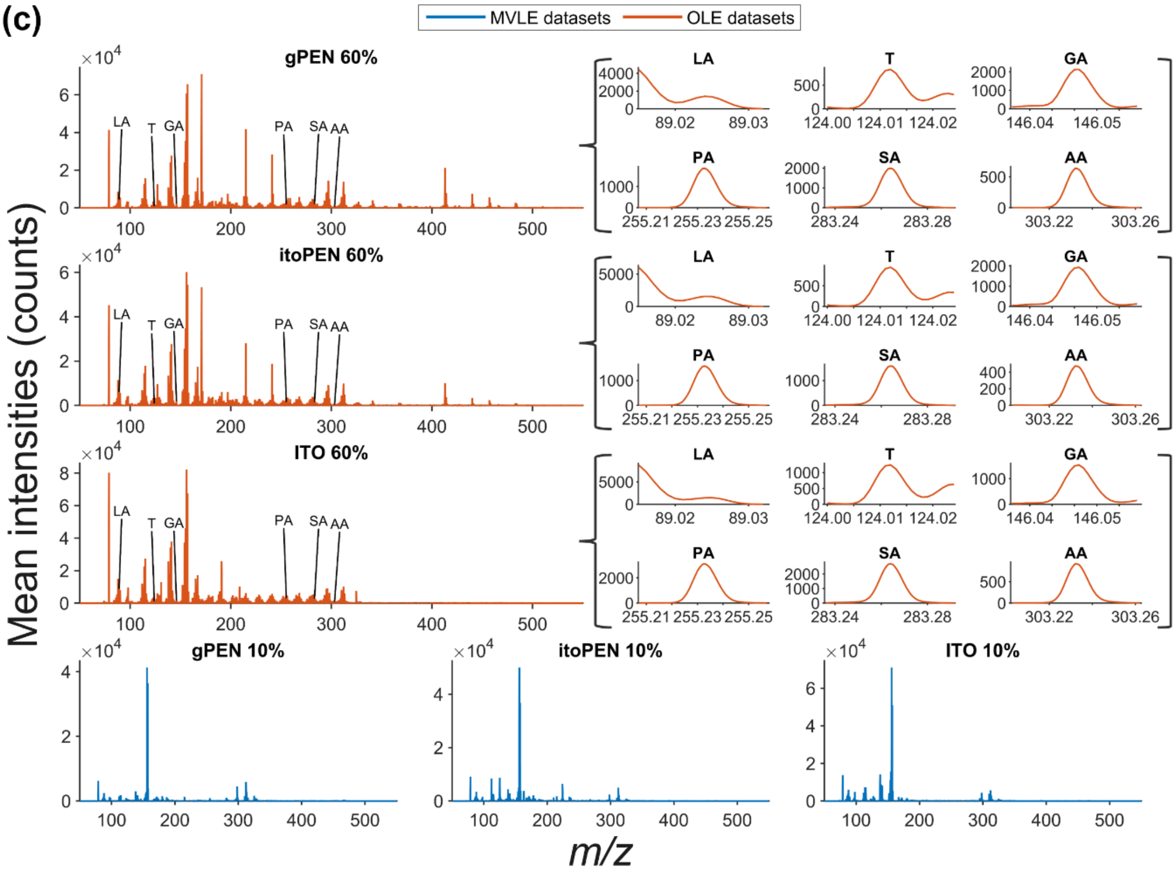
MVLE datasets
OLE datasets
gPEN 60%
itoPEN 60%
ITO 60%
gPEN 10%
itoPEN 10%
ITO 10%
LA
T
GA
PA
SA
AA
Mean intensities (counts)
m/z

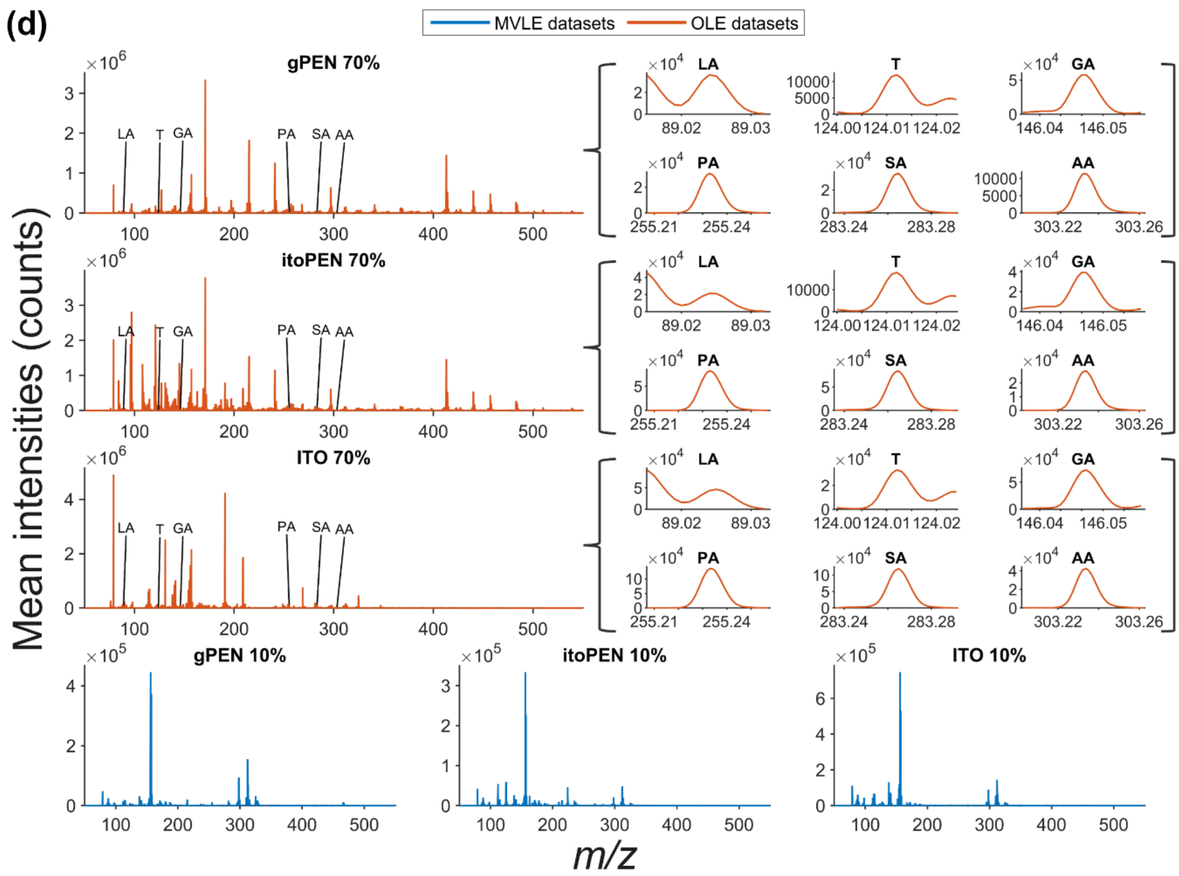


Figure S8 - On-tissue mean spectra for (a) MALDI 20 µm datasets on all substrates, (b) MALDI 5 µm datasets on all substrates, (c) MALDI-2 5 µm datasets on all substrates, (d) MALDI-2 20 µm datasets on all substrates. For all spectra, OLE spectra are displayed in orange with accompanying zoom-ins on putatively assigned peaks of ions of interest and MVLE spectra are displayed in blue.
